# Efficient construction of Markov state models for stochastic gene regulatory networks by domain decomposition

**DOI:** 10.1101/2023.11.21.568127

**Authors:** Maryam Yousefian, Anna-Simone Frank, Marcus Weber, Susanna Röblitz

## Abstract

**Background:** The dynamics of many gene regulatory networks (GRNs) is characterized by the occurrence of metastable phenotypes and stochastic phenotype switches. The chemical master equation (CME) is the most accurate description to model such stochastic dynamics, whereby the long-time dynamics of the system is encoded in the spectral properties of the CME operator. Markov State Models (MSMs) provide a general framework for analyzing and visualizing stochastic multistability and state transitions based on these spectral properties. Until now, however, this approach is either limited to low-dimensional systems or requires the use of high-performance computing facilities, thus limiting its usability.

**Results:** We present a domain decomposition approach (DDA) that approximates the CME by a stochastic rate matrix on a discretized state space and projects the multistable dynamics to a lower dimensional MSM. To approximate the CME, we decompose the state space via a Voronoi tessellation and estimate transition probabilities by using adaptive sampling strategies. We apply the robust Perron cluster analysis (PCCA+) to construct the final MSM. Measures for uncertainty quantification are incorporated. As a proof of concept, we run the algorithm on a single PC and apply it to two GRN models, one for the genetic toggle switch and one describing macrophage polarization. Our approach correctly identifies the number and location of metastable phenotypes with adequate accuracy and uncertainty bounds. We show that accuracy mainly depends on the total number of Voronoi cells, whereas uncertainty is determined by the number of sampling points.

**Conclusions:** A DDA enables the efficient computation of MSMs with quantified uncertainty. Since the algorithm is trivially parallelizable, it can be applied to larger systems, which will inevitably lead to new insights into cell-regulatory dynamics.

## Background

Within a cell, gene expression levels are governed by molecular regulators interacting with each other in gene regulatory networks (GRNs). These networks generate complex dynamical activities like oscillations or multistability that are crucial to the survival, behavior, development, and reproduction of the living cell. Hence, various mathematical models have been developed to study the dynamic behavior of GRNs [1], including Boolean models [2] and ordinary differential equation models [3]. In addition, experimental evidence has been provided that distinct cellular phenotypes correspond to stable attractors in the high-dimensional gene expression state space [4].

It has also been recognized that gene expression is a fundamentally stochastic process [5, 6], where low reactant numbers can lead to significant statistical fluctuations in molecule numbers and reaction rates. These findings have motivated studies of stochastic state-switching in gene networks. For example, stochastic changes in particular patterns of gene expression have been identified with spontaneous phenotype transitions that can diversify otherwise identical cell populations [7]. In terms of mathematical modeling, introducing stochasticity often turns originally irreversible into (weakly) reversible switches. That means particular phenotypes correspond to regions in the state space, which we call *metastable* regions or *metastabilities*, where the dynamics remains for a long time before it rapidly switch to another phenotype. These switchings are *rare events*, i.e., the dynamical system is characterized by a *separation of time scales*.

Numerous mathematical frameworks have been developed to model and analyze the dynamic behavior of stochastic GRNs dynamics, e.g., stochastic differential equations [8], or probabilistic Boolean networks [9]. The most accurate description that accounts for stochasticity due to molecular-level fluctuations and propagates dynamics according to chemical rate laws is given by the Chemical Master Equation (CME) [10, 11]. The CME is a set of ordinary differential equations (ODEs), one ODE for each possible state of the system, whereby a state is a vector containing the discrete entities of the involved species/biomolecules. It describes the time evolution of the probability density in this state space. The dimension of the CME, i.e. the number of possible system states, depends upon the total number of molecules present and the precise form of the chemical reactions and is often very large or even unbounded. Hence, the CME is usually analytically intractable except for some simplified model systems. Nevertheless, trajectories can be generated by the Stochastic Simulation Algorithm (SSA) [12]. For multistable systems, however, the simulation of phenotype switchings would require very long and/or many simulations because exploration of the state space is hindered by rare events. An example involving the dynamics of two species *A* and *B* is shown in Fig. 1. Here, simulations reveal that the system is most likely in one of the two-phase space regions corresponding to A/B being low/high or high/low, with rare transitions between these two phenotypes, whereas the phenotypes low/low and high/high do not occur.

**Figure 1.**
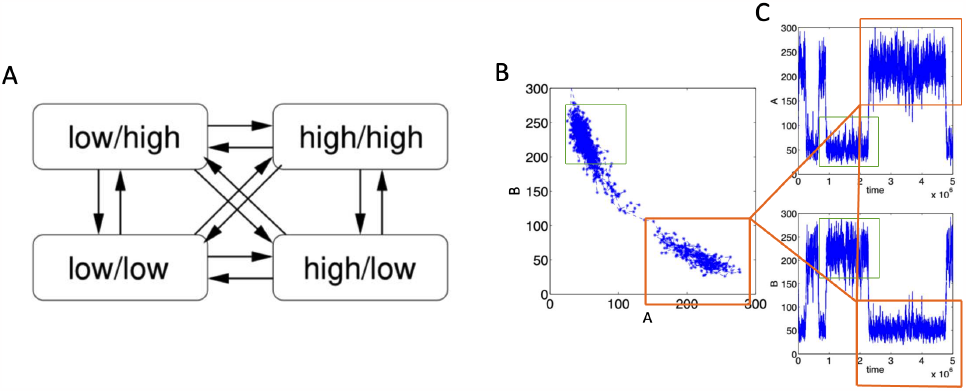
Simulation of phenotype switching. **A** The aim is to identify metastable phenotypic regions in the state space and their corresponding transition patterns. **B** These are regions in the phase space where the dynamics is most likely located. **C** Transitions between these regions are rare events and, therefore, rarely observable in stochastic simulations. For example, the region highlighted by the red rectangles corresponds to a phenotype where *A* is high while *B* is low.

Alternatively, information about phenotypes and their transition patterns can be extracted directly from analyzing the CME operator or, in case of a truncated, finite state space, from the corresponding rate matrix. In particular, the number of metastabilities, i.e. phenotypes, corresponds to the number of eigenvalues close to zero, whereas the location of the phenotypes and their transition patterns can be extracted from the corresponding eigenvectors. This information can subsequently be used to project the high-dimensional dynamical system onto the metastable sets, which results in a low-dimensional approximation of the dynamics on the slow time scales. This reduced representation of a stochastic multistable dynamical system is known as *Markov state model* (MSM) [13]. The MSM contains all information about holding times, transition rates, and transition pathways between the metastable sets (phenotypes). In a proof-of-concept study, Chu et al [14] found that MSMs provide a general framework for analyzing and visualizing stochastic multistability and state transitions in GRNs. Their study, however, was restricted to two-dimensional systems because constructing the CME rate matrix requires an enumeration of the state space, which becomes tricky in higher dimensions. In a follow-up study from the same group, Tse et al. [15] used a discretization of the state space in terms of Voronoi cells, referred to as micro-states, in combination with a sampling-based approach to obtain a finite-dimensional approximation of the CME operator, from which they derived a MSM. This approach allowed them to build a MSM for systems involving up to eight species. Even though the authors validated their results with SSA simulations, they did not provide any error estimates on the computed quantities. In addition, their approach required the use of high-performance computing (HPC) facilities, which might prevent other researchers from testing this method.

In this paper, we present an alternative and computationally more efficient approach to identifying the dynamics of cellular phenotypes from stochastic GRNs. We suggest combining the domain decomposition approach (DDA) in terms of Voronoi cells with short-time adaptive SSA simulations and statistical error estimation to calculate MSMs with quantified uncertainty. Because this method is trivially parallelizable, our approach paves the way towards constructing MSMs on single workstations with multiple CPU cores, thus making it accessible to a larger research community. We provide a proof-of-concept by applying the approach to two different networks of mutually inhibiting gene pairs with different mechanisms of self-activation. These are frequently occurring motifs in transcriptional regulatory networks to control cell fate decisions.

## Materials and methods

### The Chemical Master Equation (CME)

We consider the dynamics of *D* different species, represented by the state vector **x**(*t*) ∈ ℕ ^*D*^, under *R* prescribed reactions. Each reaction 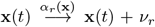 is described by a transition intensity or reaction propensity *α*_*r*_ : ℕ ^*D*^ ↦ ℝ_+_ and the stoichiometric vector *ν*_*r*_ ∈ ℕ ^*D*^. The master equation is then given by^[1]^

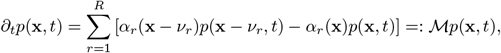

where *p*A(**x**, *t*) denotes the probability for the system to be in state **x** at time *t*. The adjoint operator ℳ^∗^, which satisfies

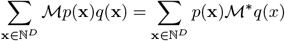

for any pair (*p, q*) of not necessarily normalized or positive functions defined over ℕ ^*D*^, is given by

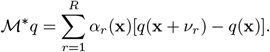

In particular, ℳ^∗^ applied to a constant function yields zero, ℳ^∗^**1** = 0.

In the following, we assume the existence of a steady state (or stationary) probability distribution *π*(*x*) satisfying ℳ*π*(**x**) = 0 ∀ **x** ∈ ℕ ^*D*^. The peaks in this stationary distribution correspond to the metastable regions that we are looking for. These are the regions in which the system is most likely to be found.

Although the CME is infinite in size, the most probable states are those near the metastable regions, and the stationary distribution tends to zero as **x** tends to infinity. Therefore, it is often reasonable to cut the system off and reduce it to some finite state space Ω ⊆ ℕ ^*D*^, thus turning the CME operator ℳ into a rate matrix. If the states **x** ∈ Ω are numbered (**x**_1_, **x**_2_, …, **x**_*N*_) such that the reactions can be written as 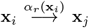, then the master operator can be written as reaction rate matrix *K* ∈ ℝ^*N* ×*N*^ with elements

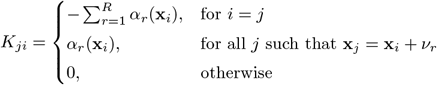

All off-diagonal elements are non-negative, and the columns sum to zero. In matrix notation, the CME reads **p**^′^(*t*) = *K***p**(*t*), where **p**(*t*) = [*p*(*x*_1_, *t*), *p*(*x*_2_, *t*), …]^⊤^. The stationary distribution ***π*** = [*π*(*x*_1_), *π*(*x*_2_), …]^⊤^ satisfies *K****π*** = 0. However, for algorithmic reasons, it might be easier to work with transition probabilities instead of transition rates.

### From transition rates to transition probabilities

By the Hille-Yosida theorem, ℳ is the infinitesimal generator of a strongly continuous semigroup 𝒯 ^(*τ*)^, and consequently the CME admits a unique continuous solution in the form *p*(*x, t* + *τ*) = 𝒯 ^(*τ*)^*p*(*x, t*). The corresponding matrix will be denoted by *T* (*τ*). *T* (*τ*) is a row stochastic transition probability matrix. For small time step *τ*, its entries are given by

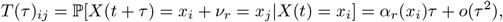

but exact analytical expressions are not available. For finite state space, *T* ^(*τ*)^ can be computed explicitly as the matrix exponential, *T* (*τ*) = exp(*τ K*^*T*^).

*T* (*τ*) can be used to compute the probability density at multiples of the lag-time *τ* via the Chapman-Kolmogorov equation,

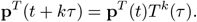

Moreover, *T* (*τ*) possesses the same eigenvectors as *K*. In particular, the stationary density satisfies ***π***^⊤^*T* (*τ*) = ***π***^⊤^.

### Partitioning into metastable regions

The CME satisfies conservation of mass, i.e. 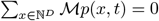. In a metastable dynamical system, this mass conservation still applies approximately to the metastable regions. In other words, the probability flux in and out of a metastable set is nearly zero. Let us now assume that there are *n*_*c*_ such metastable sets. We call a partition of unity 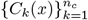 of the state space ℕ^*D*^ with 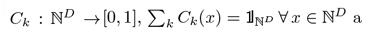 a *metastable function partitioning* if for any probability distribution *p*(*x, t*) and all functions *C*_*k*_(*x*):

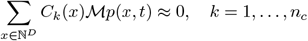

This definition can be restated in terms of the adjoint ℳ^∗^ as ℳ^∗^*C*_*k*_(*x*) ≈ 0. One possibility to search for such functions *C*_*k*_(*x*) is to reformulate the problem as an eigenvalue problem and identify metastable functions as right eigenfunctions of the adjoint operator ℳ^∗^ for eigenvalues close to zero, ℳ^∗^*C*_*k*_(*x*) = *λ*_*k*_*C*_*k*_(*x*), *λ*_*k*_ ≈ 0. Correspondingly, the discrete eigenvalue problem reads

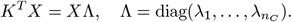

To summarize, metastability is characterized by the occurrence of a cluster of *n*_*c*_ *>* 1 eigenvalues 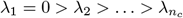 close to zero. Note that the eigenvalue *λ*_1_ = 0 is equivalent to the Perron root *λ*_1_ = 1 (also called the leading eigenvalue or dominant eigenvalue) of the corresponding *T* (*τ*). Due to this clustering of eigenvalues, the individual eigenvectors are not well-conditioned, but the subspace spanned by all leading eigenvectors is well-conditioned, and the information about the localization of the metastable regions can be extracted from this subspace via the robust Perron cluster analysis (PCCA+) [16].

### Phenotype identification using PCCA+

The aim of PCCA+ is to find a reversible matrix *A* that linearly transforms the eigenvectors *X* into *membership vectors χ* = *XA* such that *χ* forms a positive partition of unity (i.e. *χ*_*j*_(*i*) ≥ 0 ∀*i* ∈ {1, …, *N*}, *j* ∈ {1, …, *n*_*C*_} and 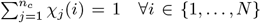. The membership vectors 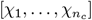 form a fuzzy clustering of the state space in that they assign to any state the probabilities for belonging to any of the *n*_*c*_ clusters, which represent the metastable regions. In general, *A* is not unique, but PCCA+ finds an optimal transformation such that the entries of the membership vectors are as close as possible to either 1 or 0. It maximizes the so-called crispness of these vectors [16]. The membership vectors not only decode the information about the localization of metastable regions but they can also be used to construct a coarse-grained representation of the GRN dynamics. This is done by projecting the dynamics onto the metastable regions by

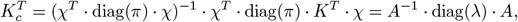

where 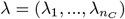. The statistical weights *w*_*k*_ of the clusters can be computed by 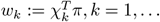, *n*_*c*_. Similarly, the normalized partial densities of the clusters are given by *π*_*k*_ := diag(*π*) *· χ*_*k*_*/w*_*k*_. While the coarse-grained transition rates might be difficult to interpret, the corresponding coarse-grained transition probabilities for a time interval of length *τ* can be obtained via the matrix exponential

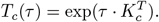

This reduced representation of the dynamics as *T* (*τ*) of a low-dimensional Markov chain is referred to as MSM. The coarse-grained matrices preserve the dominant eigenvalues and, hence, the time scale of the slow processes. Hence, propagation of any projected density vector *χ*^*T*^ **p** commutes with the projection of the propagated density *K***p**,

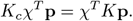

The bottleneck of this approach is the size of the matrix *K* (or *T* (*τ*), respectively). The matrix size increases exponentially with the number of species. Even if truncated to a reasonable finite state space size, and even though *K* is generally sparse, the dimension might still be too large for algorithms based on matrix multiplications, like the one for computing the eigenvalues and eigenvectors. One solution is to decompose the state space ℕ ^*D*^ into a finite number of relatively small states, which corresponds to a Galerkin discretization of the CME operator.

### Discretization

Given a partition of unity 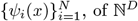, the metastable functions *C*_*k*_(*x*) can be approximated by

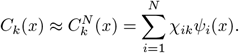

The condition Σ_*k*_ *χ*_*ik*_ = 1 is sufficient to ensure that 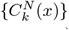 is a partition of unity if 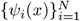 is one. Hence, the vectors *χ*_*k*_ = (*χ*_1*k*_, …, *χ*_*Nk*_)^*T*^ can be interpreted as *membership vectors* whereby the entry *χ*_*ik*_ ∈ [0, 1] indicates the membership of function *ψ*_*i*_(*x*) to the metastable function *C*_*k*_(*x*).

Using ℳ^∗^*C*_*k*_(*x*) = *λ*_*k*_*C*_*k*_(*x*), *λ*_*k*_ ≈ 0 and the approximated *C*_*k*_(*x*), and forming inner products^[2]^ with test functions 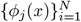 we obtain the matrix eigenvalue problem,

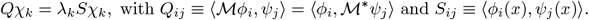

Thus, the vectors *χ*_*k*_ can be computed as right eigenvectors of the matrix pair (*Q, S*) corresponding to eigenvalues close to zero. Since the state space is high-dimensional, we must use meshless ansatz functions. One possibility is Voronoi cells. Assume we choose the test functions equal to the ansatz function. Then *S*_*ij*_ ≡ *δ*_*ij*_ and

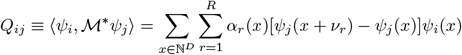

The evaluation of this sum would require the detection of all states *x* in the support of *ψ*_*i*_ from where the support of *ψ*_*j*_ can be reached within one reaction step, which is very difficult or even impossible on unstructured Voronoi tesselations in high dimensions. Alternatively, we can work with transition probabilities. A Galerkin discretization of 𝒯 ^(*τ*)^ in terms of Voronoi cells 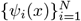 and test functions *ϕ*_*i*_(*x*) = *ψ*_*i*_(*x*)*π*(*x*) ≡ *π*_*i*_(*x*) results in the discrete eigenvalue problem *Pχ*_*k*_ = *λ*_*k*_*χ*_*k*_ with

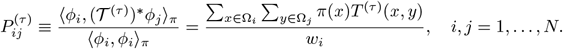

If we have sampled points 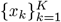 according to the partial stationary density *π*_*i*_(*x*), then *P*_*ij*_ can be approximated by importance sampling as

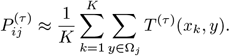

Even though we do not know the stationary density point-wise, we can still sample it by running stochastic simulations with SSA. Hence, we present the following algorithm for constructing a MSM.

### Algorithmic Approach

Fig. 2 illustrates the main steps of the proposed algorithm, which we will elaborate on in the subsequent subsections a)-f). The algorithmic parameters are explained in Tab. 1.

**Figure 2.**
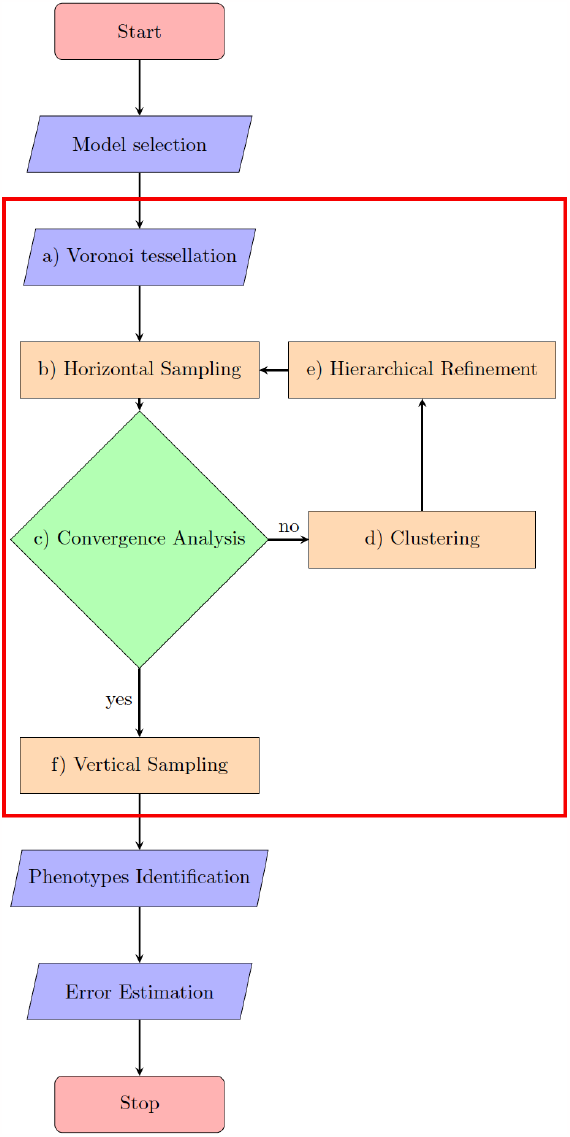
Algorithmic workflow for the current study, starting with the selection of the network to be analyzed within the framework: **a** The state space is decomposed into a finite number of cells represented by their center nodes. **b** The stationary density is sampled within each cell (horizontal sampling). **c** convergence towards the stationary density is checked. If convergence occurs, **f** points are propagated forward over time (vertical sampling). Otherwise, **d**,**e** the horizontal sampling points are clustered to find the location of two new center nodes, and the cell is refined hierarchically. Horizontal sampling and refinement are repeated for the refined cells until convergence is reached. The algorithmic steps within the red box constitute the proposed DDA. Afterward, the PCCA+ algorithm is used to identify metastable regions (phenotypes), and the uncertainty in the model output is estimated.

#### a) Voronoi tessellation

Given the potentially high-dimensional nature of the state space, we employ meshless methods, particularly Voronoi tessellations. A Voronoi tessellation with center points {*x*_1_, …, *x*_*N*_ } ∈ ℕ ^*D*^ decomposes the state space ℕ^*D*^ into a number of *N*_0_ cells. Each cell is then described by its characteristic function *ϕ*_*i*_(**x**) : ℕ^*D*^ → {0, 1}, *i* = 1, …, *N*_0_ with

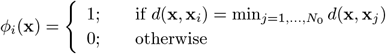

where *d*(**x, x**_*i*_) is a suitable distance function between **x** and **x**_*i*_, in our case the Euclidean metric. Note that this decomposition of the state space does not require an a priori truncation of the state space to some finite set Ω.

#### b) Horizontal sampling

The aim of this step is to sample locally from the stationary distribution within each Voronoi cell. For this purpose, we initiate multiple SSA trajectories in the center point of each Voronoi cell. If a trajectory tries to leave the cell, the next state proposed in the SSA algorithm is rejected, and the trajectory stays in its current state until the Markov process jumps to another state within the cell. For each trajectory, we keep track of the time when it first tries to exit the cell. The average of these times over all trajectories that were initiated within the same cell serves as an estimate for the mean first exit time from this cell. In addition, a comparison of the different sampling chains within one cell is used to check for convergence towards the local stationary distribution.

#### c) Convergence analysis

A necessary criterion for checking the convergence of multiple sampling chains towards the same distribution is the Gelman-Rubin convergence indicator [17]. The criterion compares the within-chain-variances with the between-chain-variances of different trajectories and computes a *potential scale reduction factor* (*R*), which should ideally be close to one [17]. We compute *R* after *N*_hor_ horizontal sampling points. If it is still above a defined threshold, namely, 1+*R*_thr_, then the sampling did not converge towards its equilibrium distribution. Thus, the horizontal trajectories are continued with *N*_hor_ more sampling points and *R* is computed again. As soon as *R* falls below the threshold, the horizontal sampling is stopped. Our assumption is that, in this case, the horizontal sampling has reached an equilibrium. If this does not happen within rep_max_ *· N*_hor_ sampling points, then the basis function is marked for refinement because the non-convergence of the sampling indicated that there is a metastability inside the corresponding cell that has to be resolved by subdividing the cell. This is usually the case when the local stationary density has multiple maxima within a cell, e.g. when it is located on the boundary between two metastable regions. All cells marked for refinement are then ordered according to decreasing mean first exit time, and only the top ref_max_ cells are finally refined on the next level. The final goal is to refine the state space in such a way that further discretizations of the transition matrix would not change the spectral information (of the dominant eigenvalues and eigenvectors) anymore. This is the case if the cells either have a high exit rate or the sampling is rapidly mixing within the cells. Rapid mixing would lead to rapid convergence according to the Gelman-Rubin criterion [18].

#### d, e) Clustering and hierarchical refinement

The aim of this step is to decompose the cells in which the horizontal sampling did not converge into two new cells. To find the centers of the new cells, we apply *k-means* clustering to all horizontal sampling points and use the cluster centers as the center points of the new Voronoi cells. We refine the cells in a hierarchical manner. That means, if a cell with characteristic function *ϕ*_*i*_(**x**) is decomposed into two new cells *ϕ*_*i*1_(**x**) and *ϕ*_*i*2_(**x**) with center points **x**_*i*1_ and **x**_*i*2_, then the characteristic function of *ϕ*_*i*1_ is defined as

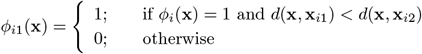

and correspondingly for *ϕ*_*i*2_(**x**). For the refined cells, the algorithm proceeds with horizontal sampling as described in step b). To effectively track the hierarchy of cells, we use a table for bookkeeping the indexes of parental nodes and their respective child nodes. The major advantage of hierarchical refinement is that the sampling only needs to be repeated for the refined cells while the already converged horizontal samplings of the remaining cells are not affected.

#### f) Vertical sampling

The aim of this step is to approximate the entries of *T* (*τ*). For this purpose, *N*_vert_ points are randomly sampled from the horizontal sampling points within each cell and propagated forward by SSA over a time interval of length *τ*. For each endpoint located in cell *j*, a transition count of one is added to the *j* entry of a transition count vector, which is finally normalized to obtain a transition probability vector **p**_*i*_. This step is illustrated in Fig. 3. The vertical sampling is repeated *N*_r_ times for each cell *i*, resulting in a set *D*_*i*_ = {**p**_*i*1_, …, **p**_*i*_*N*_*r*_ } of *N*_*r*_ possible transition probability vectors for row *P* ^*τ*^ (*i*, :). From this set, we compute the maximum likelihood estimate

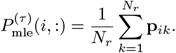

This estimate is then used to compute a MSM via PCCA+. Note that *τ* should be chosen large enough to avoid some individual cells having very low transition probabilities to other cells, as this might have a negative impact on the accuracy of the estimated MSM. On the other hand, *τ* should be as small as possible to save computing time. As a rule of thumb, we use a value of *τ* that equals approximately twice the maximum mean first exit time of all cells. As such, *τ* could be determined during the simulation run after the horizontal sampling, but we decided to use a fixed value of *τ* in each example in order to make the results comparable in case other algorithmic parameters are modified. The major advantage of this sampling-based approach is that we can control the quality of the discretization by checking the convergence behavior of the sampling and controlling the accuracy of the constructed MSM by using re-sampling techniques.

**Figure 3.**
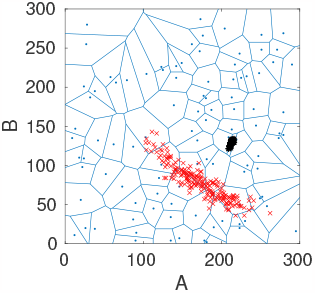
Two-dimensional Voronoi diagram (blue) with 100 cells including the points from horizontal sampling within one specific cell (black) and their end points resulting from the vertical sampling (red).

### Error estimation

While the maximum likelihood estimate of *P* ^(*τ*)^ provides a point estimate for the mean, it does not convey information regarding uncertainties in these transition probabilities. To address this issue and to quantify the size of the error matrix, it is imperative to estimate the covariance matrix [19]. To choose between the many different possible estimators, we assume that the distribution of probability vectors **p**_*i*_ follows a Dirichlet distribution. The Dirichlet distribution, Dir(*α*), comprises a group of continuous multivariate probability distributions parameterized by a vector *α* of positive real numbers. Given a set *D*_*i*_ = {**p**_*i*1_, …, **p**_*i*_*N*_*r*_ } of probability vectors, the classical way of estimating Dirichlet distribution parameters is to maximize the log-likelihood log ℙ(*D*|*α*). This is done by a Newton iteration described in [20] and implemented in the MATLAB toolboxes FASTFIT [21] and LIGHTSPEED [22]. Once an estimate of the Dirichlet parameter vector *α* is available for each row of *P* ^(*τ*)^, we sample a number of *N*_*MSM*_ transition probability matrices and compute the corresponding MSM. This allows us to visualize the uncertainty in computed quantities of interest, e.g., eigenvalues or holding probabilities, in terms of boxplots.

### Implementation of illustrative examples

All the implementations are done in MATLAB-R2021b, and the codes are available in https://github.com/sroeblitz/MSM2CME. Table 1 summarizes all algorithmic parameters and their values in the different simulation runs. All computations have been performed on a laptop with Intel Core i7-8650U CPU @ 1.90 GHz and 8 MB cache. The random seed in the Matlab code was fixed, and the same for all simulations to ensure the reproducibility of results. We apply our algorithm to two different networks of mutually inhibiting species with different mechanisms of self-activation, as illustrated in Figs. 4 and 5. The example networks were chosen because they nicely illustrate the strengths and weaknesses of our proposed algorithm while at the same time being computationally tractable on a laptop without making use of parallelization.

**Table 1:** Algorithmic parameters and their values in different simulation scenarios (A1, A2, A3, and B).

| parameter | explanation | A1 | A2 | A3 | B |
| --- | --- | --- | --- | --- | --- |
| $N_0$ | initial number of Voronoi cells | 100 | 100 | **20** | 100 |
| $N_x, N_y$ | size of the domain $[0, N_x] \times [0, N_y]$ in which the initial Voronoi cells are placed | 300 | 300 | 300 | 100 |
| $N_{\text{hor}}$ | number of horizontal sampling points until first convergence check | 500 | 500 | 500 | 500 |
| $N_{\text{traj}}$ | number of horizontal trajectories within each cell | 5 | 5 | 5 | 5 |
| $\text{rep}_{\text{max}}$ | maximum number of horizontal samplings before final convergence check | 5 | 5 | 5 | 5 |
| $R_{\text{thr}}$ | threshold $1 + R_{\text{thr}}$ for Gelman-Rubin convergence criterion | 0.03 | 0.03 | 0.03 | 0.03 |
| $\text{ref}_{\text{max}}$ | maximum number of cells to be refined on one level | 10 | 10 | 10 | 10 |
| $\text{level}_{\text{max}}$ | maximum hierarchy level | 3 | 3 | **5** | 3 |
| $N_{\text{vert}}$ | number of vertical sampling points for each cell | **100** | 500 | 500 | 200 |
| $\tau$ | lag time of the $T(\tau)$ | 5000 | 5000 | 5000 | 10 |
| $N_r$ | number of candidate rows for each cell | 10 | 10 | 10 | 10 |
| $N_{\text{MSM}}$ | number of sampled transition probability matrices for visualizing uncertainties | 20 | 20 | 20 | 20 |

**Figure 4.**
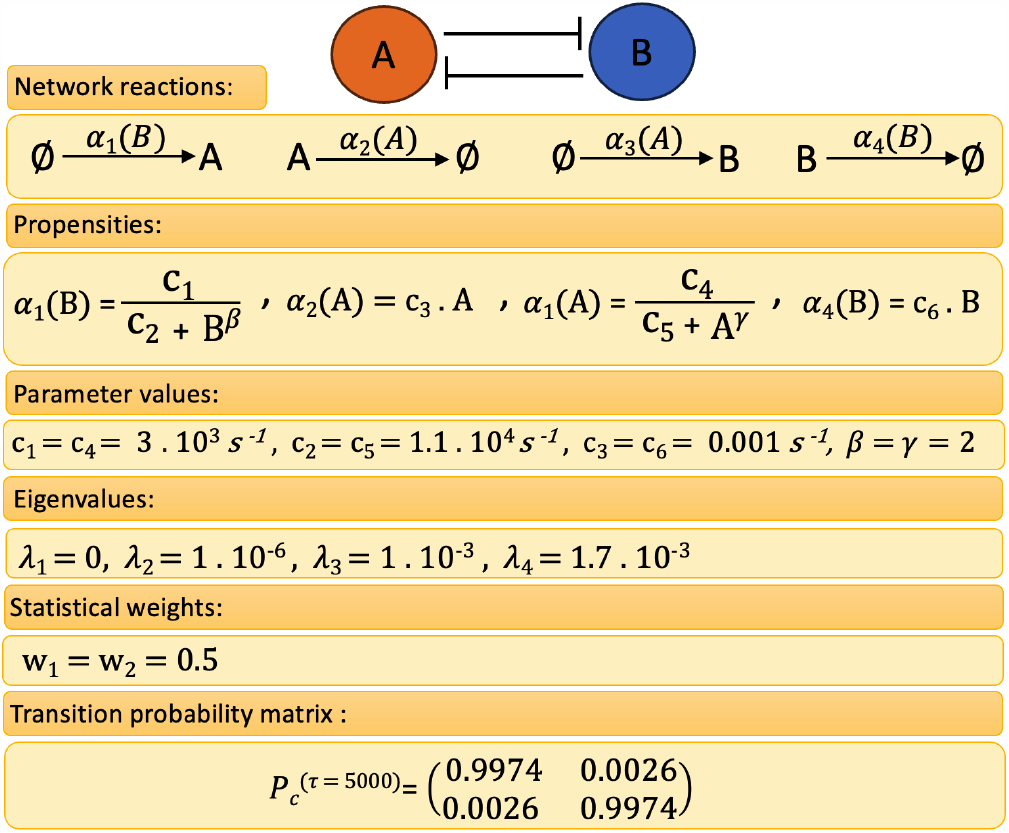
**Model A**: Representation of the toggle switch network without self-activation including network reactions, propensities, parameter values, eigenvalues, statistical weights, and transition probability matrix with lag time *τ* = 5000.

**Figure 5.**
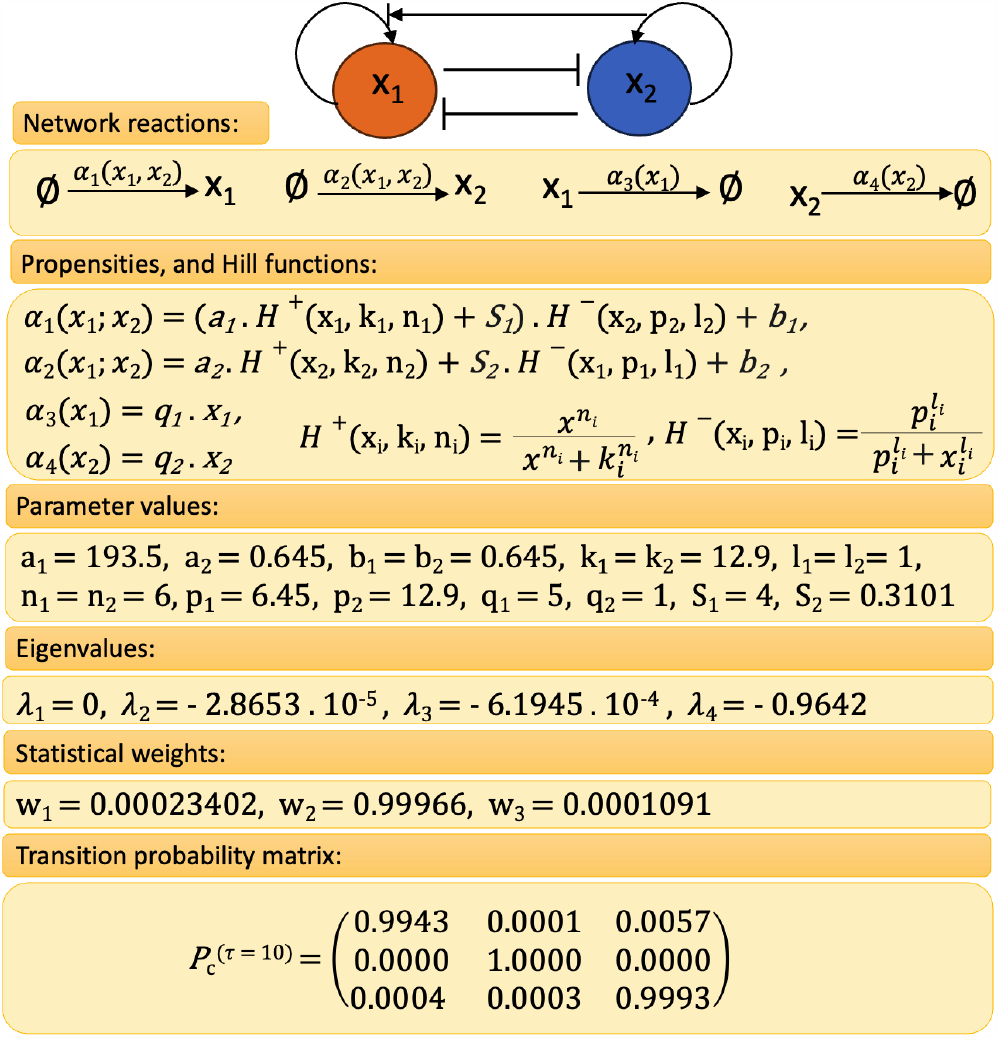
**Model B**: Representation of the macrophage polarization network including network reactions, propensities, parameter values, eigenvalues, statistical weights, and transition probability matrix with lag time *τ* = 10.

## Results

### Model A: Toggle switch

The toggle switch is composed of two repressors, further called A and B, and two constitutive promoters. Each promoter is inhibited by the repressor that is transcribed by the opposing promoter. We describe the mutual inhibition of the two genes using a stochastic, highly simplified two-stage model of gene expression based on the deterministic model presented in [23]. This model includes the synthesis and degradation of the two repressors as well as parameter values that preserve the symmetry in the system’s topology.

In Fig. 4, the toggle switch and its reactions, propensities, parameter values, eigenvalues, statistical weights, and transition probability matrix with *τ* = 5000 are shown. Spectral analysis of the full CME matrix on the domain [0, 300] *×* [0, 300] reveals a cluster of two eigenvalues close to zero, followed by two eigenvalues further away from zero (Fig. 4). The coarse-grained *P* ^(*τ*)^ for *τ* = 5000 serves as a reference solution to which we compare the MSM obtained with the DDA. Application of PCCA+ for *n*_*c*_ = 2 clusters results in two equally weighted membership functions which separate the state space along the diagonal *A* = *B* (Figs. 6b and 6c).

**Figure 6.**
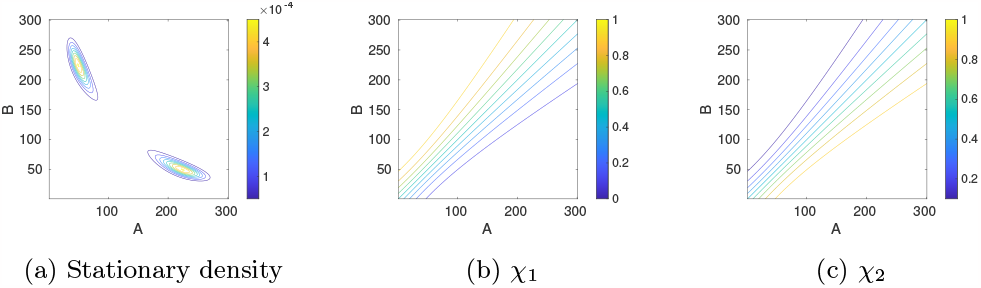
**Model A**: Reference solution on the truncated state space Ω = [0, 300] *×* [0, 300] including the stationary density as well as the two cluster membership functions *χ*_1,2_.

#### Case A1: Discretization and local sampling reveal the metastabilities

To illustrate our approach, we start with an initial discretization into 100 Voronoi cells and initialize 5 trajectories in each cell. The Gelman-Rubin convergence estimator is computed every 500 sampling steps, whereby the maximum length of the horizontal samplings is set to 5 *·* 500 steps. At most 10 cells are marked for refinement, and the maximum refinement level is 3. Fig. 7a shows how the adaptive sampling automatically refines the Voronoi cells placed in the support of the stationary density as well as in the transition region between the two metastable regions, resulting in a total of 120 Voronoi cells on the last level. For the vertical sampling, we propagate from each cell 100 points picked randomly from the horizontal sampling points over a lag time *τ* = 5000. This vertical sampling is repeated 10 times, resulting in each cell in 10 candidate probability vectors from which the parameter vector *α* of the Dirichlet distribution is estimated for the corresponding row. The overall computing time for this setting is about 25 minutes, whereby the exact time depends on the selected random seed. We then applied PCCA+ to the maximum likelihood estimate of 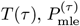, to construct an estimate of the MSM. This analysis reveals two cluster (*λ*_2_ = 0.9950) with weights *w*_1_ = 0.48386, *w*_2_ = 0.51614 and transition probability matrix

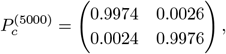

which approximates the reference MSM quite well. The two membership functions *χ*_1,2_ are plotted in Figs. 7c and 7d. As in the reference solution, they clearly divide the phase space along the diagonal (Fig. 7b). Moreover, the location of the partial stationary densities, *π*_1_ and *π*_2_, in Figs. 7e and 7f agrees well with the location of the density in the reference solution (Fig. 6a). However, the question remains whether the inaccuracies in comparison to the reference MSM are due to the discretization error, i.e. the error made in the approximation of membership functions by Voronoi cells, or due to a sampling error in approximating the entries of the transition probability matrix *P* ^(*τ*)^. To find an answer to this question, we repeat the numerical experiment with a different number of Voronoi cells and a different number of vertical sampling points.

**Figure 7.**
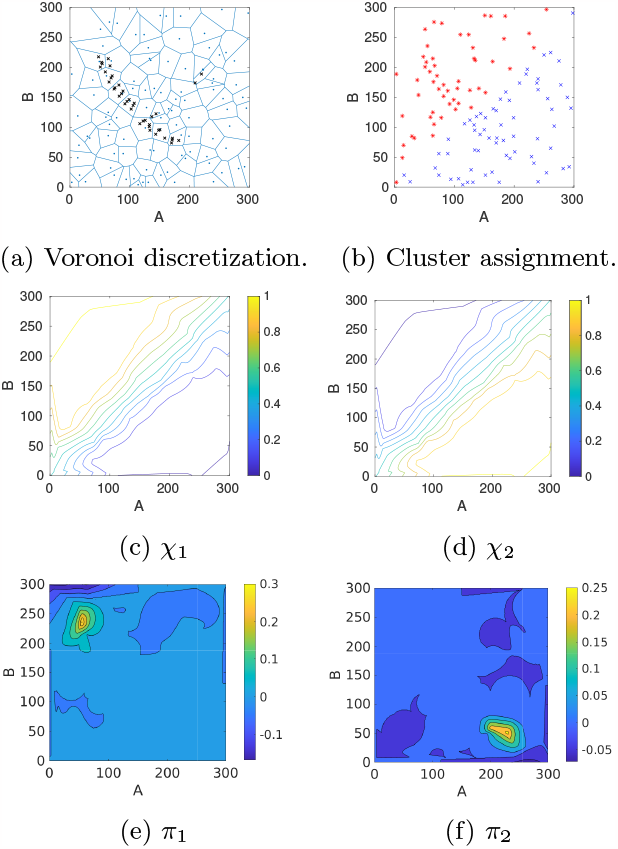
**Model A (case A1)**: **(a)** For an initial start discretization with *N*_0_ = 100 randomly located Voronoi cells (blue lines), the adaptive sampling and refinement strategy automatically places new nodes (black crosses) along the support of the stationary density as well as in the transition region between the two metastable regions. **(b)** Based on the approximated transition probability matrix, the individual cells, represented by their center nodes, are assigned to one of the two basins of attraction (blue or red). **c)**,**d)** This assignment is based on the values of the two approximated membership functions *χ*_1,2_: Nodes were colored according to the cluster to which they belong with the highest membership. **e)**,**f)**: The projected partial stationary densities *π*_1_ and *π*_2_.

#### Case A2: Increasing the number of vertical sampling points reduces the uncertainty

We repeat the numerical experiment from the previous section, but this time we increase the number of vertical sampling points per cell to *N*_vert_ = 500. All remaining algorithmic parameters remain the same. The overall computing time increases to about 2 hours. Again, by applying PCCA+ to the maximum likelihood estimate 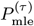, we obtain two clusters (*λ*_2_ = 0.9947) with weights *w*_1_ = 0.49246, *w*_2_ = 0.50754 and transition probability matrix

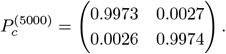

Thus, the perfect symmetry from the reference solution is much better recovered now. While there is no difference visible by eye in the plotted membership functions and partial densities, the decrease in uncertainty becomes visible when multiple MSMs are constructed from sampled transition probability matrices. The boxplots in Fig. 8 (middle column) show that the uncertainty in estimated quantities like the second eigenvalue, holding probabilities 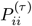, or cluster weights *w*_1,2_ significantly decreases when the number of vertical sampling points is increased.

**Figure 8.**
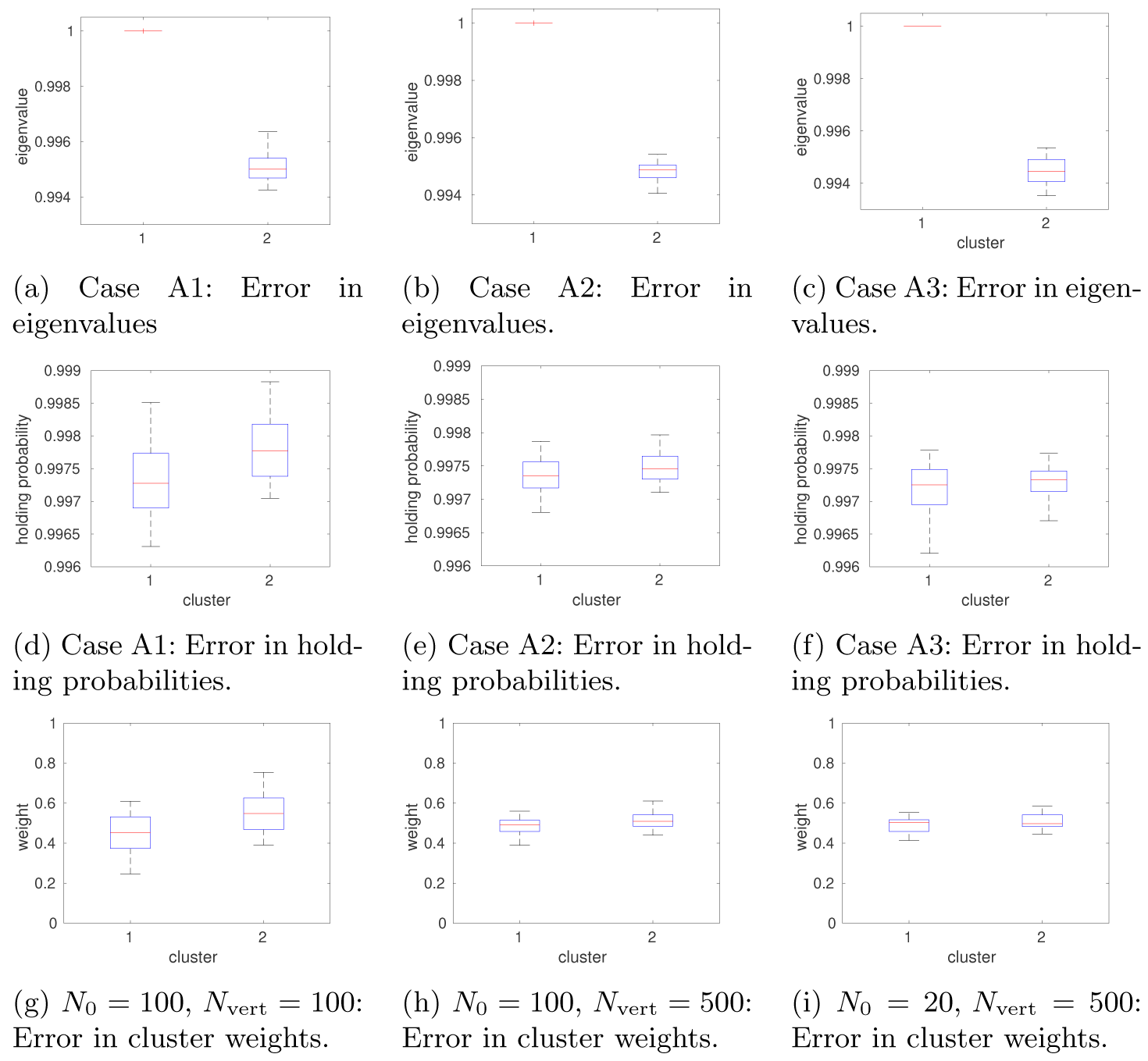
**Model A**: Increasing the number of vertical sampling points from *N*_vert_ = 100 (left) to *N*_vert_ = 500 (middle) while maintaining the same number of initial Voronoi cells (*N*_0_ = 100) and horizontal sampling points (*N*_vert_ = 500) reduces the uncertainty in the computed quantities of interest. Decreasing the number of initial Voronoi cells, *N*_0_, from 100 to 20 (right) does not deteriorate this gain in accuracy.

#### Case A3: Decreasing the number of Voronoi cells does not deteriorate the discretization error

To examine the influence of the discretization error, we keep the number of vertical sampling points at *N*_vert_ = 500 but decrease the number of initial Voronoi cells to *N*_0_ = 20 while increasing the maximum hierarchy level to level_max_ = 5 to enable sufficient refinement. All remaining algorithmic parameters remain the same. The hierarchical refinement results in a total of *N* = 60 Voronoi cells, and the overall computing time is about 1 hour. The second eigenvalues of 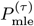 is *λ*_2_ = 0.9946. The two clusters have weights *w*_1_ = 0.5036, *w*_2_ = 0.4964 and transition probabilities

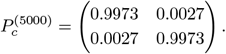

The two membership functions *χ*_1,2_ are plotted in Figs. 9c and 9d. They still divide the phase space approximately along the diagonal (Fig. 9b), though not as symmetric as in the reference solution because the second cluster *χ*_2_ (high A, low B) stretches slightly into the region above the diagonal. However, since this mainly happens in the region where the stationary density is low, it does not affect the symmetry of the dynamics. The estimated quantities of interest are nearly equally close to the reference solution as in the case with *N*_0_ = 100, and the uncertainties are only slightly larger (Figs. 8c, 8f and 8i). However, the discretization error in the approximation of the partial stationary densities *π*_1_ and *π*_2_ (Figs. 9e and 9f) is larger with fewer Voronoi cells (compare Fig. 6a).

**Figure 9.**
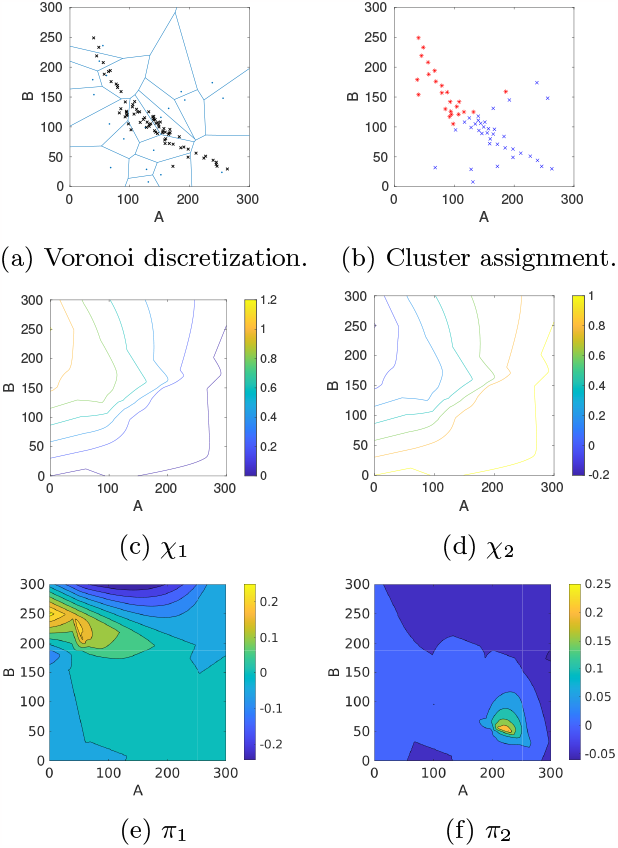
**Model A (case A3)**: **(a)** For an initial start discretization with *N*_0_ = 20 randomly located Voronoi cells (blue lines), the adaptive sampling and refinement strategy automatically places new nodes (black crosses) along the support of the stationary density as well as in the transition region between the two metastable regions. **(b)** The individual cells, represented by their center nodes, are assigned to one of the two basins of attraction (blue or red). To simplify visualization, nodes have been colored according to the cluster to which they belong with the highest membership. **c)**,**d)** This assignment is based on the values of the two approximated membership functions *χ*_1,2_: Nodes are colored according to the cluster to which they belong with the highest membership. **e)**,**f)** The projected partial stationary densities *π*_1_ and *π*_2_ illustrate the location of the metastable regions.

### Model B: Macrophage polarization

The macrophage polarization model presented in [24] consists of four elementary reactions corresponding to synthesis and degradation of STAT1 (*x*_1_) and STAT6 (*x*_2_), respectively, as well as four stochastic propensity functions presented in Fig. 5. We consider a parameter set from [24] for which the system has three metastable regions corresponding to STAT1/STAT6 being low/high, low/low, or high/low, as visualized by the potential energy landscape *U* (*x*) = − log(*π*(*x*)) in Fig. 10a. Spectral analysis of the full CME matrix on the domain [0, 100]*×*[0, 100] reveals a cluster of three eigenvalues close to zero followed by a gap to the fourth smallest eigenvalue. The parameter values, eigenvalues, and statistical weights are listed in Fig. 5. Application of PCCA+ to the corresponding eigenvectors results in a decomposition of the state space in terms of three membership functions *χ*_1,2,3_ as illustrated in Figs. 11a-11c with statistical weights and transition probability matrix shown in Fig. 5. All three clusters are clearly metastable, but most of the statistical weight is on cluster 2 (low/high), which also has the largest basin of attraction. This makes it very difficult to detect the other two attractors in SSA simulations that are initiated outside their basins of attractions. In contrast, our domain decomposition approach with *N*_0_ = 100 cells and *N*_vert_ = 200 sampling points clearly identifies the three metastable regions in terms of their corresponding membership functions (Figs. 10 and 11). During the horizontal sampling, the algorithm particularly refines cells located at the boundary of cluster 1 as well as cells close to the support of the stationary density *π*(*x*), resulting in a total of 117 cells on the last refinement level 3 (Fig. 10b). The computing time for this example was about 35 min.

**Figure 10.**
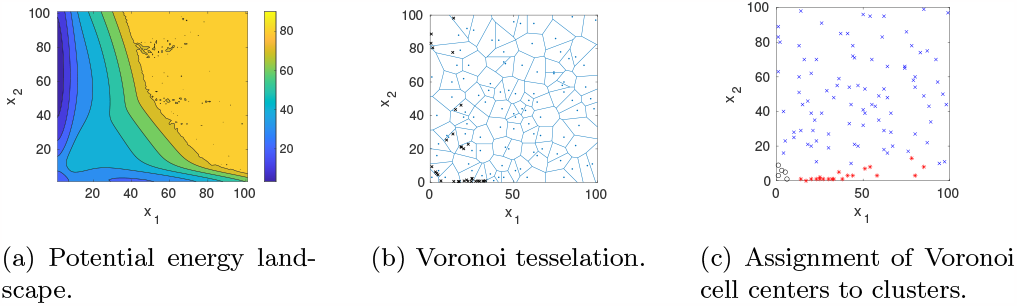
**Model B**: **(a)** The potential energy landscape *U* (*x*) = − log(*π*(*x*)) constructed from the full CME clearly shows three different basins of attraction corresponding to *x*_1_*/x*_2_ being low/high, low/low or high/low. **(b)** The adaptive sampling and refinement places new cells at the boundaries of the local stationary densities and in the transition regions. **(c)** The clustering algorithm assigns each Voronoi cell center to one of the three metastable regions.

**Figure 11.**
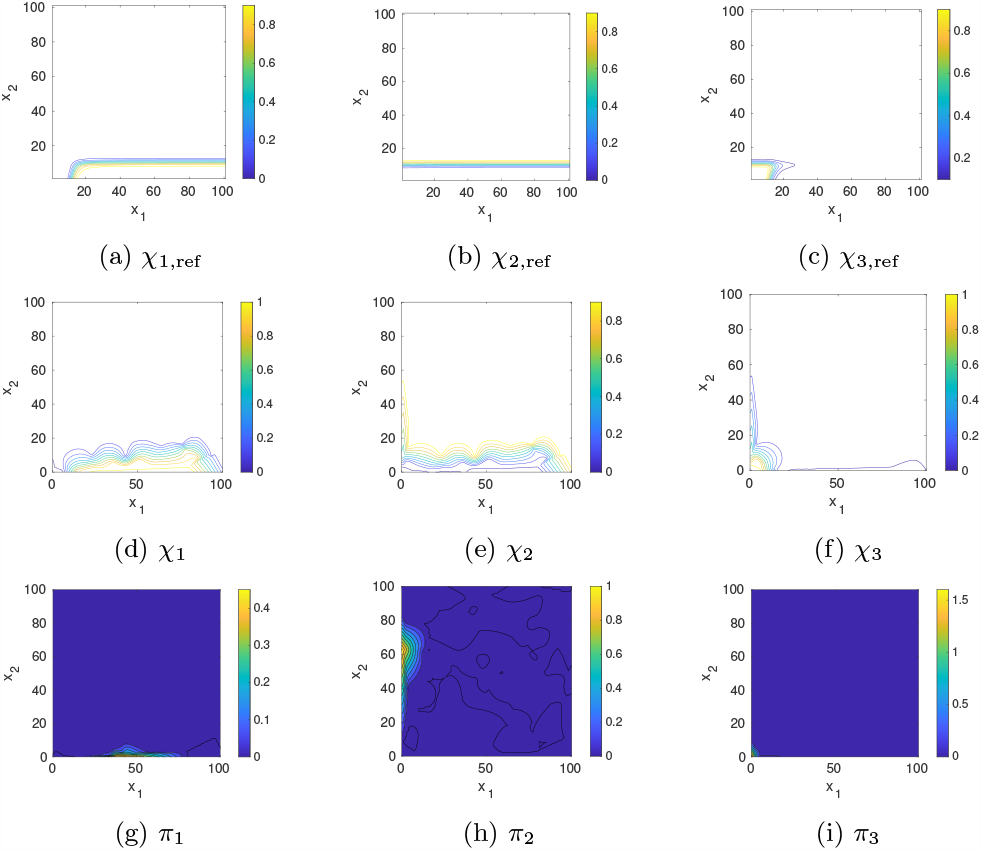
**Model B**: Contour plot of the corresponding membership functions *χ*_1,2,3_ computed from the full CME matrix (top) and their approximation by Voronoi cells (middle) as well as the approximated partial stationary densities (bottom).

The maximum likelihood estimate 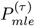 has a cluster of three eigenvalues close to one (*λ*_1_ = 1, *λ*_2_ = 0.9993, *λ*_3_ = 0.9944), and PCCA+ returns three clusters with weights *w*_1_ = 0.0038, *w*_2_ = 0.0043, *w*_3_ = 0.9918 and transition probability matrix

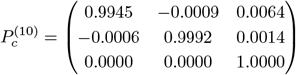

Note that negative entries in this matrix can occur due to the membership vectors not being perfectly zero or one [16]. However, these negative entries are small. The major difference to the reference solution is in the statistical weights. Since the exit probability from cluster 3 is approximated as being almost zero, its statistical weight is close to one, in contrast to the reference solution, where cluster 2 gets the highest statistical weight. When sampling multiple MSMs, cluster 3 is assigned the lowest statistical weight in almost all scenarios, in some cases cluster 1 has the highest weight, whereas cluster 2 never dominates (Fig. 12c). Increasing the number of vertical sampling points to *N*_vert_ = 500 was not sufficient to improve the approximation of the weights. Note that if the number of initial Voronoi cells is decreased, e.g. to *N*_0_ = 20, it might even happen that none of the initial Voronoi centers is placed in the basin of attraction of the low/low cluster, in which case it could not be detected by the algorithm (Fig. 13).

**Figure 12.**
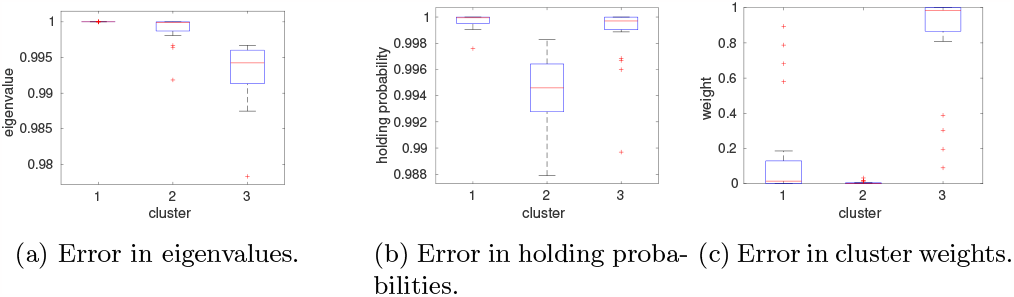
**Model B**: The eigenvalues and holding probabilities are approximated with reasonably small uncertainty, while the cluster weights cannot be inferred with sufficiently small uncertainty.

**Figure 13.**
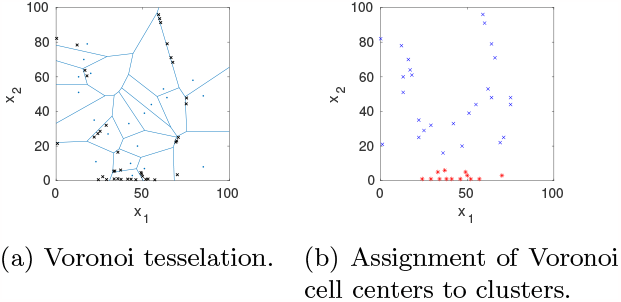
**Model B**: If none of the cell centers is placed in the lower left corner, i.e. in the basin of attraction for the cluster where both *x*_1_ and *x*_2_ are small, then this metastable region is not detected by the algorithm.

## Discussion

Our algorithm provides a tool for the theoretical examination of stochastic state-switching within gene networks. The results show that the proposed domain decomposition approach is capable to discover all metastable regions in stochastic gene regulatory networks with multiple attractors and to approximate the membership functions with good accuracy. A metastable region is found as long as at least one initial center node is located in its basin of attraction. If the basins of attraction are large enough, this usually happens even with randomly placed initial cells. However, metastable regions with a very small basin of attraction might be missed in this approach, as we have illustrated for model B.

The algorithm also succeeds in approximating the partial stationary densities within the identified metastable regions as well as their global weights, as illustrated for model A. However, the approximation of the global statistical weights fails if the clusters are dynamically well separated. This is not surprising as the stationary density of an almost decoupled Markov chain is known to be highly ill-conditioned [25, 26]. However, once the partial stationary densities have been located, there exist alternative methods to correctly infer transition rates between them, for example transition path sampling [27, 28]. Another solution would be to increase the lag time *τ* in our algorithm, which would also increase the overall computing time. Typically, the horizontal sampling is fast whereas the vertical sampling takes most of the computing time. This time scales linearly with both the number of vertical sampling points, *N*_vert_, the number of candidate rows, *N*_*r*_, and the lag time *τ*.

In general, the accuracy of the approximated MSM depends mainly on the number of Voronoi cells, whereas the uncertainty is largely influenced by the number of vertical sampling points. Thus, if the main point of interest is to explore the number and location of metastable phenotypes, the algorithm can be run quickly with a low number of vertical sampling points. Since the CME is only a model itself, the modelling error might dominate the approximation error of the MSM. Still, understanding the characteristics of metastable regions and the barriers separating them provides critical insights into the stability and transition dynamics of gene expression profiles. In particular, elucidating the critical points where stochastic state-switching events are more likely to occur, further deepens our comprehension of the regulatory mechanisms governing gene expression variability.

Although we have implemented the algorithm for only two-dimensional systems so far, the use of a meshless discretization makes it easily extendable to higher dimensional systems and other biological applications where multistability is expected to occur.

Our future work will be to parallelize the algorithm. Since both the horizontal as well as the vertical samplings are independent between different cells, the proposed algorithm is trivially parallelizable in that the samplings for different cells can be run in parallel. In addition, different horizontal sampling chains within one cell could run in parallel as well. This possibility for parallelization paves the way towards the application to higher dimensional problems, though high performance computing facilities might be needed for systems with a large accessible state space.

## Conclusion

We have proposed an algorithm for the computation of MSMs, which is based on local adaptive samplings and the statistical assembly of these trajectories into a global transition probability matrix. In our approach, many local short-term simulations can be combined to figure out the long-term global behavior of the system. The method is also adaptive such that computational costs are saved, and sampling data is only generated where it is “needed”. Our algorithm identifies basins of attraction of metastable regions as well as the local stationary densities with moderate computational effort. It also succeeds in approximating their statistical weights and transition probabilities as long as the lag time of the Markov state model is chosen large enough to capture sufficiently many transitions between the metastable regions. Our algorithm is trivially parallelizable, which paves the way for its application to larger systems.

## Ethics approval and consent to participate

Not applicable.

## Consent for publication

Not applicable.

## Availability of data and materials

The datasets generated and/or analysed during the current study are available in the github repository https://github.com/sroeblitz/MSM2CME.

## Author’s contributions

MY wrote the first draft of the manuscript, created the software, and analyzed the simulation results. ASF reviewed and edited the manuscript. MW developed the methodology, reviewed and edited the manuscript. SR acquired the funding, conceptualized the study, developed the methodology, created the software, analyzed the simulation results, reviewed and edited the manuscript.

## Acknowledgment

Not applicable.

## Footnotes

[1] We postulate that *α*_*r*_ (**x**) = 0 if ∃*i* : *x*_*i*_ *<* 0 or ∃*i* : *x*_*i*_ + (*ν*_*r*_)_*i*_ *<* 0.

[2] We consider inner products in the weighted *l*^2^ sequence space, 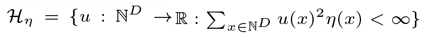 with 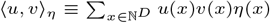 In the unweighted sequence space where 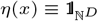, we simply omit *η* in the notation.

